## Supplemental Figures for "Disruption of myelin structure and oligodendrocyte maturation in a pigtail macaque model of congenital Zika infection"

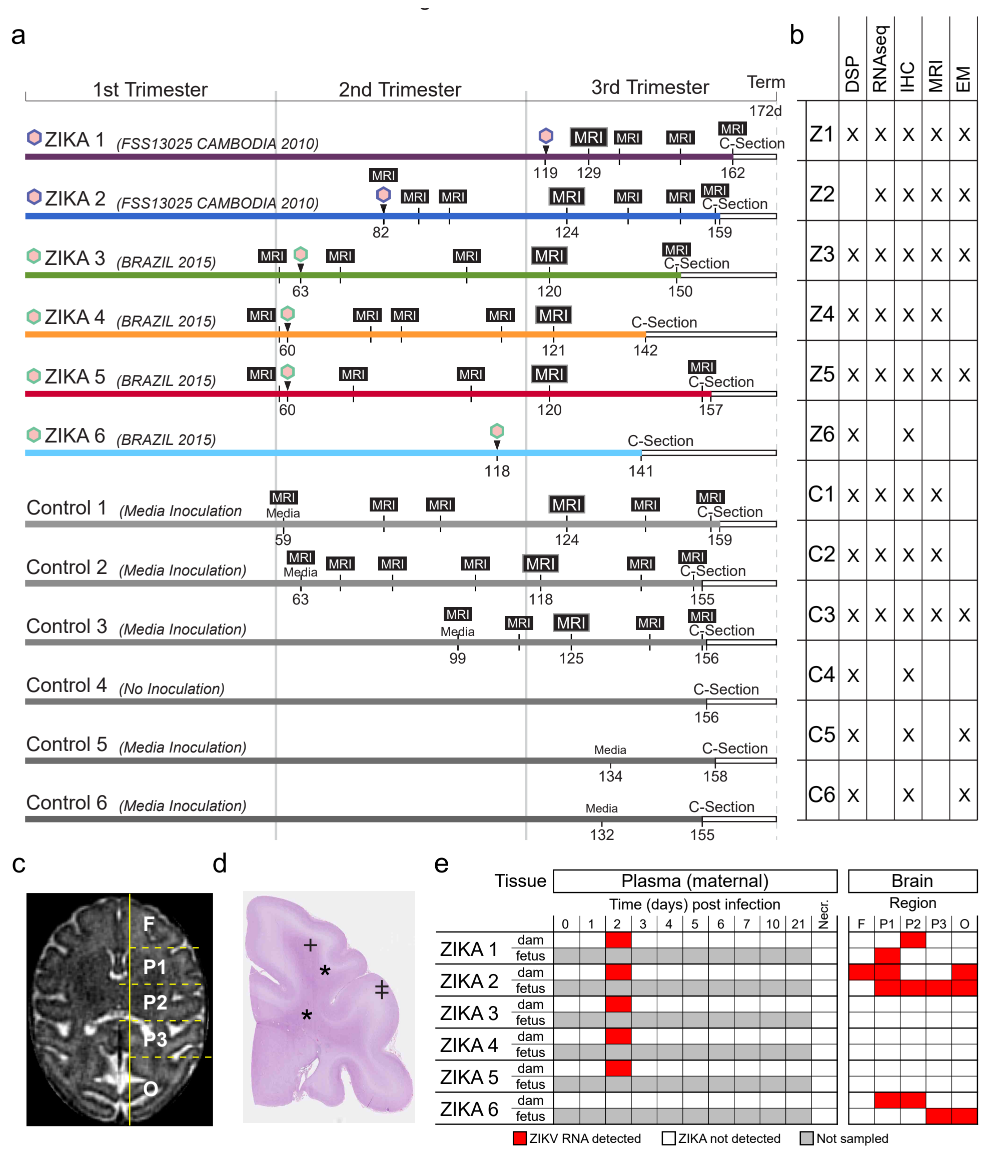


**Figure S1. Maternal ZikV infection study design and NHP fetal brain sampling overview. a**) A timeline of study events for ZIKA1-6 and CTL1-6 is shown with respect to the gestational age in days on the x-axis. Pregnant dams were inoculated subcutaneously with Asian-lineage ZikV clinical isolate from Cambodia (FSS13025 Cambodia 2010; GenBank no. MH368551) or American-lineage ZikV virus clinical isolate from 2015 Brazil (Brazil 2015; GenBank no. KX811222) at the gestational age indicated. Dams underwent Cesarean section at the indicated gestation age, prior to 172 days, which is the average gestational age at spontaneous delivery in the Washington National Primate Research Center (WaNPRC) colony. Magnetic resonance imaging (MRI) was performed at indicated time points. **b**) Table representing animals from which tissue was used in downstream analysis. For each assay, tissue was included for analysis only if it matched the brain location of other sections being analyzed in the same assay. This led to some brain areas for some animals not being represented in the final dataset. **c)** Fetal cerebrum bulk tissue dissection scheme overlaid onto an MRI of a control animal at 156 gestational days (GD) in the coronal plane. The cerebrum was sectioned at the midline (parasagittal vertical line) and coronal sections (transverse dashed lines) collected from 5 regions, referred to as frontal (F), parietal (P1-P3), and occipital (O). Tissue from right hemisphere was used fresh for bulk RNA sequencing or preserved in formalin or 4% paraformaldehyde for immunohistochemical and electron microscopy (EM) analyses. **d**) The left hemisphere was submerged in formalin or 4% paraformaldehyde (PFA) and coronal slices, corresponding to P2 and O, were sectioned and embedded into paraffin for immunohistochemical (IHC) staining. The representative micrograph is a hematoxylin and eosin (H&E)-stained section from the parietotemporal region (P2) of a control animal at 155 GD. EM samples were collected from cerebral gray and white matter (*). Tissue samples used for bulk RNA sequencing were collected in the superficial grey matter (‡). Quantification of protein expression was performed in the superior gyri (+) of the parietal lobe where myelin was most dense. **e)** Table representing the detection of ZikV RNA by TaqMan polymerase chain reaction on tissue from dam (upper row) and fetus (lower row) for each ZikV-exposed animal. Left table, plasma samples; right table, brain samples.

Figure S2

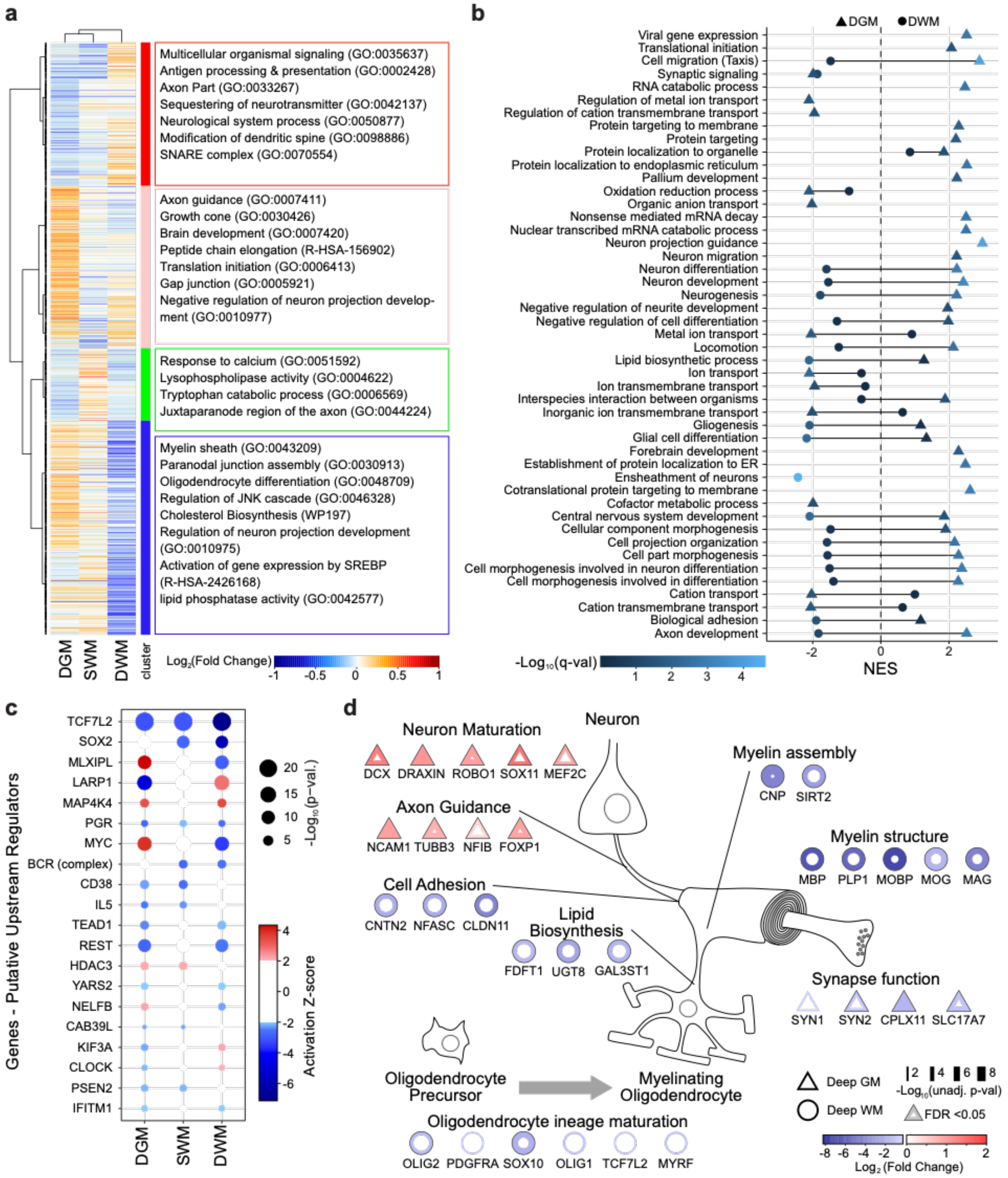


**Figure S2. Spatial transcriptomic analysis of NHP fetal brain responses to ZikV infection in discrete brain regions. a**) **Left,** Heatmap of average $\log_2$ fold change by ROI of 1234 differentially expressed genes identified as significant (FDR<0.05) in at least one brain region (**Table S3**). Orange indicates upregulated gene expression in ZikV relative to CTL. Blue indicates down-regulated gene expression in ZikV relative to CTL. Hierarchical clustering using Euclidean distance measure and ward.D2 identified 4 clusters of genes (vertical color bar). **Right**, functional pathway identification by Over Representation Analysis (ORA) is displayed for each cluster, according to color (**Table S4**). **b**) Plot of normalized enrichment scores (NES) for pathways identified by GSEA in DGM (triangles) and DWM (circles) regions based on analysis of all genes detected. Color scale indicates adjusted q-values (**Table S5**). **c**) Dot plot of 20 significant upstream transcriptional regulators (TR) identified in at least two out of the three brain regions (DGM, SWM, and DWM) of interest using Ingenuity Pathway Analysis (IPA; **Table S6**). IPA Upstream Regulator analysis was performed on the 1234 DEG identified in panel a (**Table S3**). The dot size is proportional to significance (-log10 *p*-value); color represents the TR activation z-score (> 2 for activation or < -2 for inhibition). Genes were sorted by sum of absolute z-scores across the three ROIs, and the top three genes, *TCF7L2*, *SOX2*, and *MLXIPL*, have the most statistically significant overall z-scores. **d**) Functional schematic showing cellular location of genes involved in neuron health and synaptic function linked with oligodendrocyte development and myelination. Genes are colored by log fold change expression (ZikV *vs*. CTL) identified in the DSP analysis. Shape indicates the region represented in the analysis, with DGM in triangles and DWM in circles. Line thickness signifies the negative log of the unadjusted p-value, and the presence of a thin bounding black line indicates FDR<0.05.

Figure S3

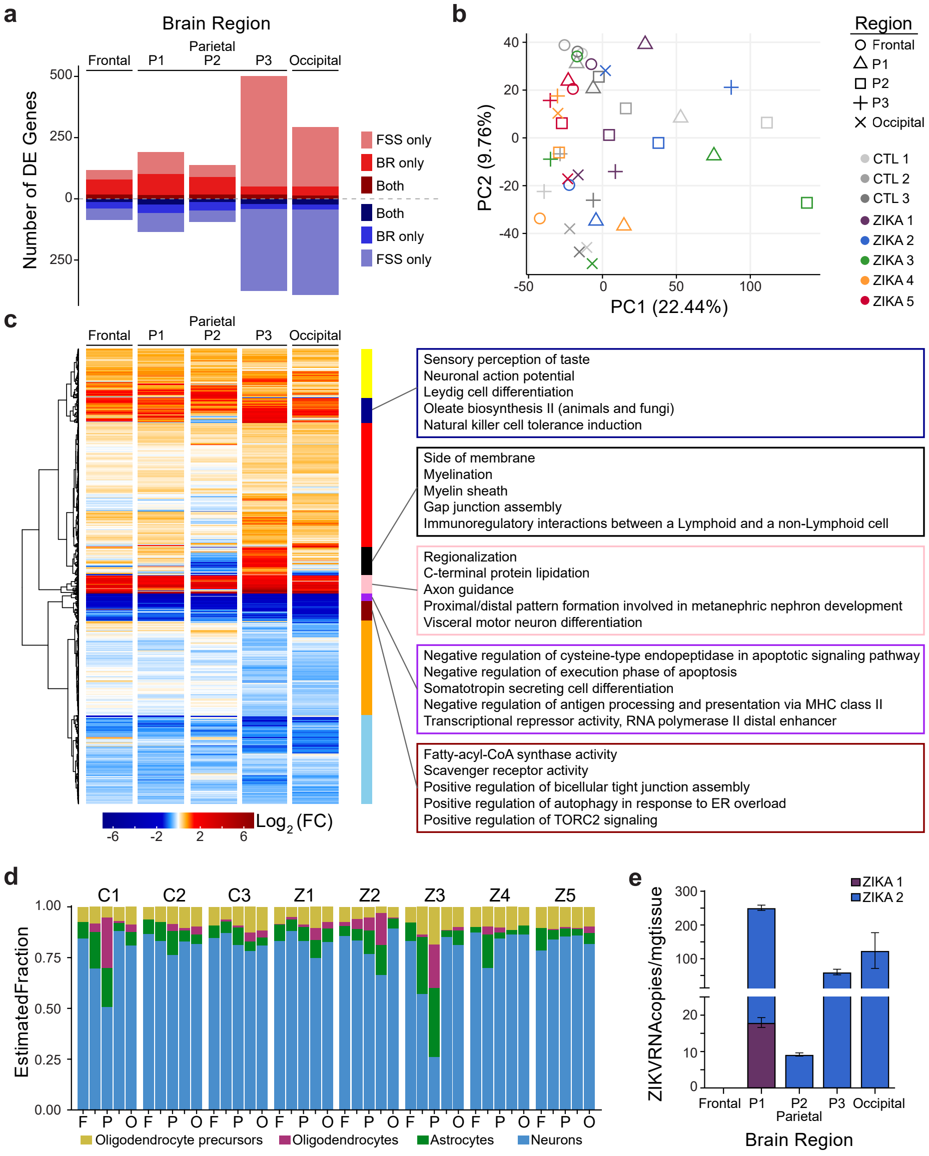


**Figure S3. Global RNA sequencing of ZikV and control NHP fetal superficial cortex. a**) Bar plot of the total number of differentially expressed genes (DEG) identified in comparisons of ZikV *vs.* control (CTL) for brain region-matched samples. Red indicates up-regulated genes; blue indicates down-regulated genes. The color shade represents the number of DEG in ZikV/FSS, ZikV/BR or both strains. **b**) Principal component analysis (PCA) of the whole transcriptome across all brain samples. Each point in the plot represents an individual sample, with brain region denoted by symbol and animal denoted by color. **c**) Heatmap of average log fold change (LFC) expression of 1505 DEG identified in at least one brain region ($p<0.05$). Hierarchical clustering was performed using Euclidean distance measure and ward.D2 and identified 9 clusters of genes (vertical color bar) (**Table S7**). Right, functional pathway identification by over representation analysis against gene ontology, KEGG and wikipathways pathway databases is displayed for the five clusters with the largest average differences in gene expression, according to color (**Table S8**). **d**) Stacked bar chart of relative percent abundance of individual cell types predicted in each brain sample using deconvolution analysis with CIBERSORT (**Table S9**). Bars are ordered left to right from frontal (F), parietal (P)1-3, and occipital (O) regions within each animal. Animal is listed across the top. Bar color denotes cell type, including oligodendrocyte precursors (gold), oligodendrocytes (purple), astrocytes (green), and neurons (blue). **e**) ZikV RNA was detected in the brain of two animals using a ZikV-specific qRT-PCR assay. Maternal ZIKA1 and ZIKA2 animals were inoculated with ZikV/FSS and fetal brain samples analyzed at 43 and 77 days post-inoculation, respectively (see **Figure S1;** note that ZikV RNA was detected in fetal brain of ZIKA6 using a different primer set and therefore not included in the quantitative representation in panel **S3e**). Average gestational age (±SD) of ZikV-exposed vs CTL animals in RNAseq analysis=157(±2) vs 154(±8) days; p=0.6 by t-test.

Figure S4

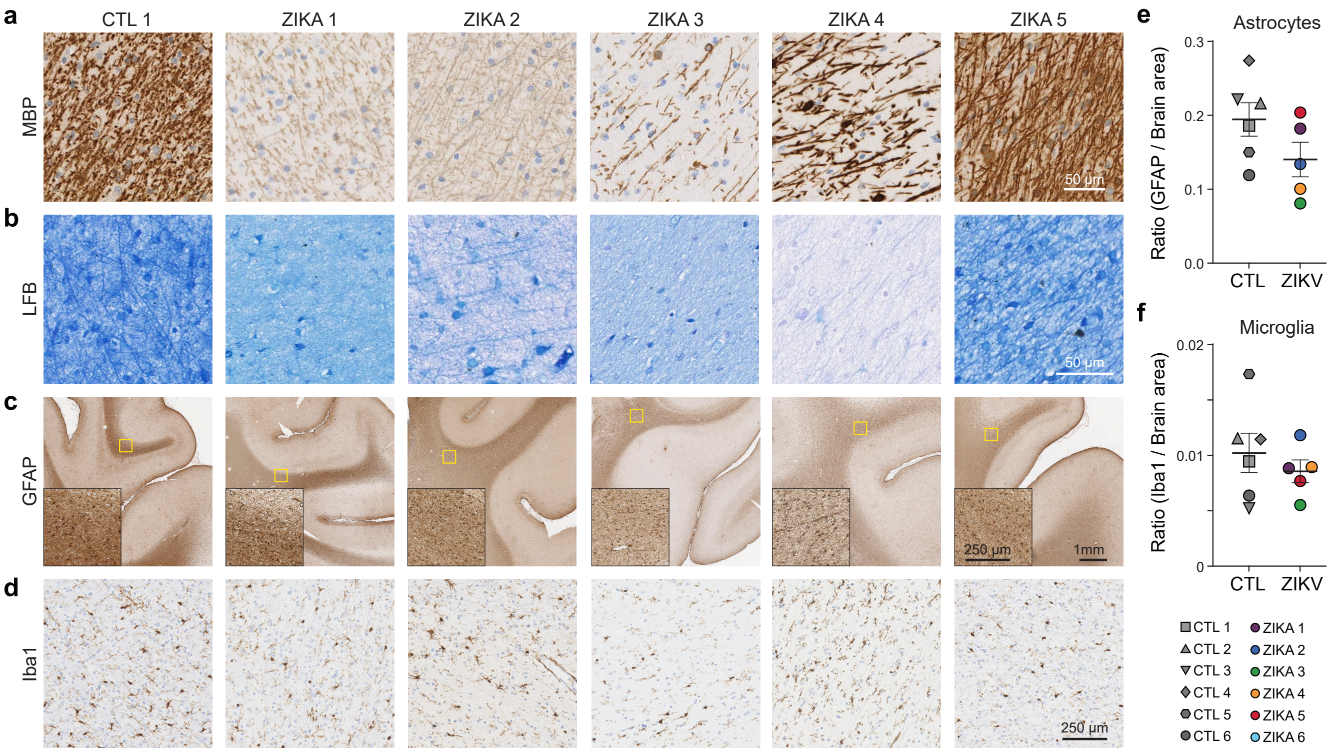


**Figure S4. Immunohistological characterization of ZikV and control NHP fetal brains.** Formalin-fixed paraffin-embedded (FFPE) coronal slices of parietal cortex were used for immunohistochemical (IHC) analyses. **a**) High-magnification micrographs of myelin basic protein (MBP) IHC from central areas of deep white matter (DWM) containing dense fiber tracts, demonstrating reduced intensity of MBP signal, as well as decreased density of MBP-labeled fibers in ZikV virus cohort animals. **b**) Luxol fast blue combined with periodic acid-Schiff (LFB-PAS) staining was used to detect compact myelin. **c**) IHC for astrocyte marker, glial fibrillary acidic protein (GFAP), with insets demonstrating high-magnification areas of the DWM tracts. **d**) IHC for the microglial marker, allograft inflammatory factor 1 (AIF-1/Iba), obtained from the DWM tracts. The regions selected are approximately the same location as the inset images in panel **c**. **e-f**) Quantification of **e**) GFAP and **f**) Iba1 signal throughout the parietal cortex expressed as the ratio of area occupied by the chromogen divided by the total area of brain tissue present within the section. A single section per animal was quantified. Points in the plots represent individual animals, with bars indicating mean and standard error mean (SEM). Differences between ZikV and CTL animals were not significant for either GFAP or Iba1. Average gestational age (±SD) of ZikV-exposed vs CTL animals in IHC analysis=152(±2) vs 154(±8) days; $p=0.47$ by t-test.

### Figure S5

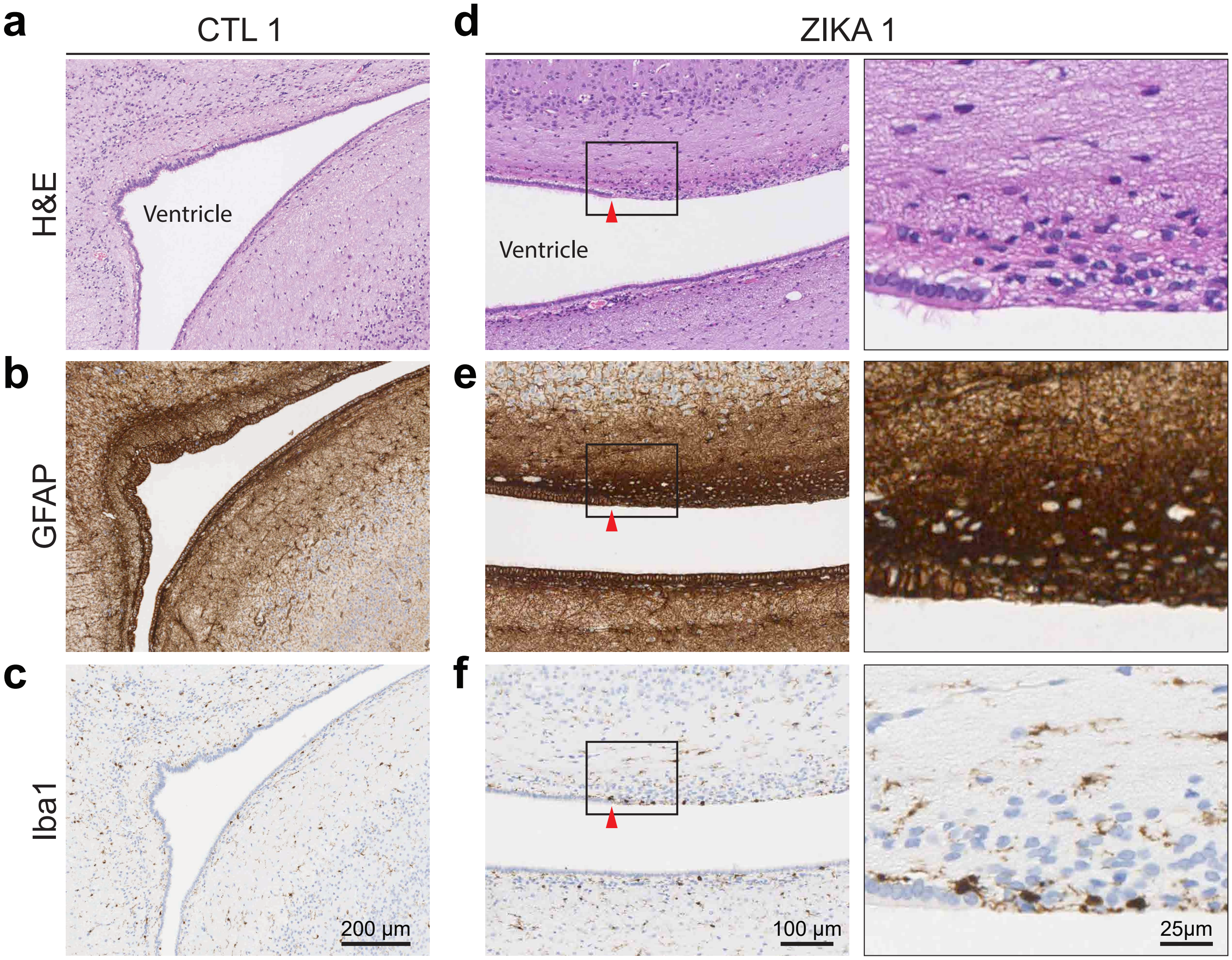


**Figure S5. Histopathologic changes in the periventricular region of the occipital cortex lesion of ZikV and control NHP fetal brain.** Formalin-fixed paraffin-embedded (FFPE) coronal section of occipital cortex was used for hematoxylin and eosin (H&E) staining and immunohistochemical (IHC) analyses. **a-c**) H&E, glial fibrillary acidic protein (GFAP) and allograft inflammatory factor 1 (AIF-1/Iba1) stained sections of the lateral ventricle and ependymal lining of control (CTL1) for reference. **d-f**) Similar staining of the same region in a ZikV virus cohort animal (ZIKA1), with inset in the right column of images showing high-magnification of the area marked by the black rectangle. Red arrowheads indicate a transition point where the ciliated ependymal lining shows an area with multiple vacuoles and increased cellularity. This area has increased GFAP staining intensity and a higher density of Iba1-positive microglia. Qualitatively similar findings were observed in ZIKA3 and ZIKA4 animals.

Figure S6

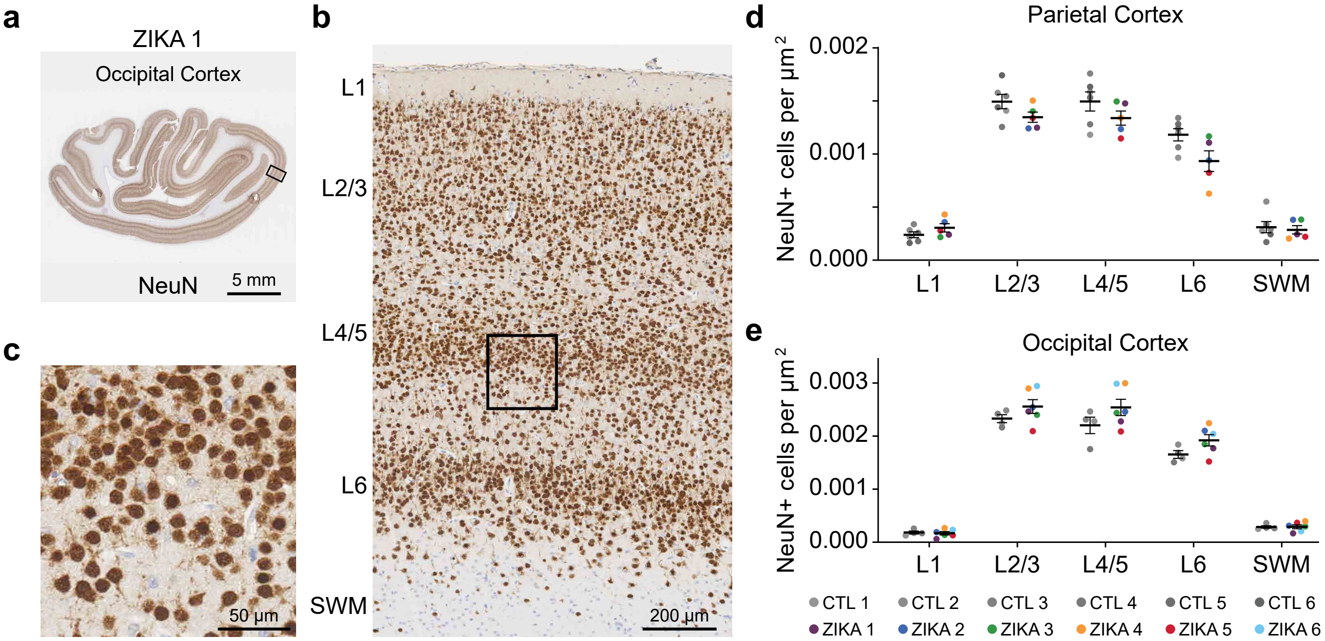


**Figure S6. Neuron density by cortical layer in parietal and occipital cortex of ZikV and control NHP fetal brains.** Formalin-fixed paraffin-embedded (FFPE) coronal sections of parietal and occipital cortex were used for immunohistochemical (IHC) analyses. **a-c**) Neuronal nuclear protein (NeuN)-stained coronal section of occipital cortex in a ZikV virus cohort animal (ZIKA 1). **b**, cortical layers 1-6 (L1-L6) and subcortical white matter (SWM) are indicated. **c**) inset shown by the black rectangle in panel b of the neuronal bodies. **d-e**) Semi-automated quantification of neuron density in the **d**) parietal and **e**) occipital cortex. Regions of interest (ROI) delineating individual cortical layers (L1, L2/3, L4/5, L6) and SWM were manually drawn, as in panel b, and NeuN+ cells were counted within each ROI using automated Visiopharm software. Points in the plots represent individual animals, from which three independent areas containing all cortical layers were quantified and averaged together, with bars indicating mean and standard error mean (SEM).

### Figure S7

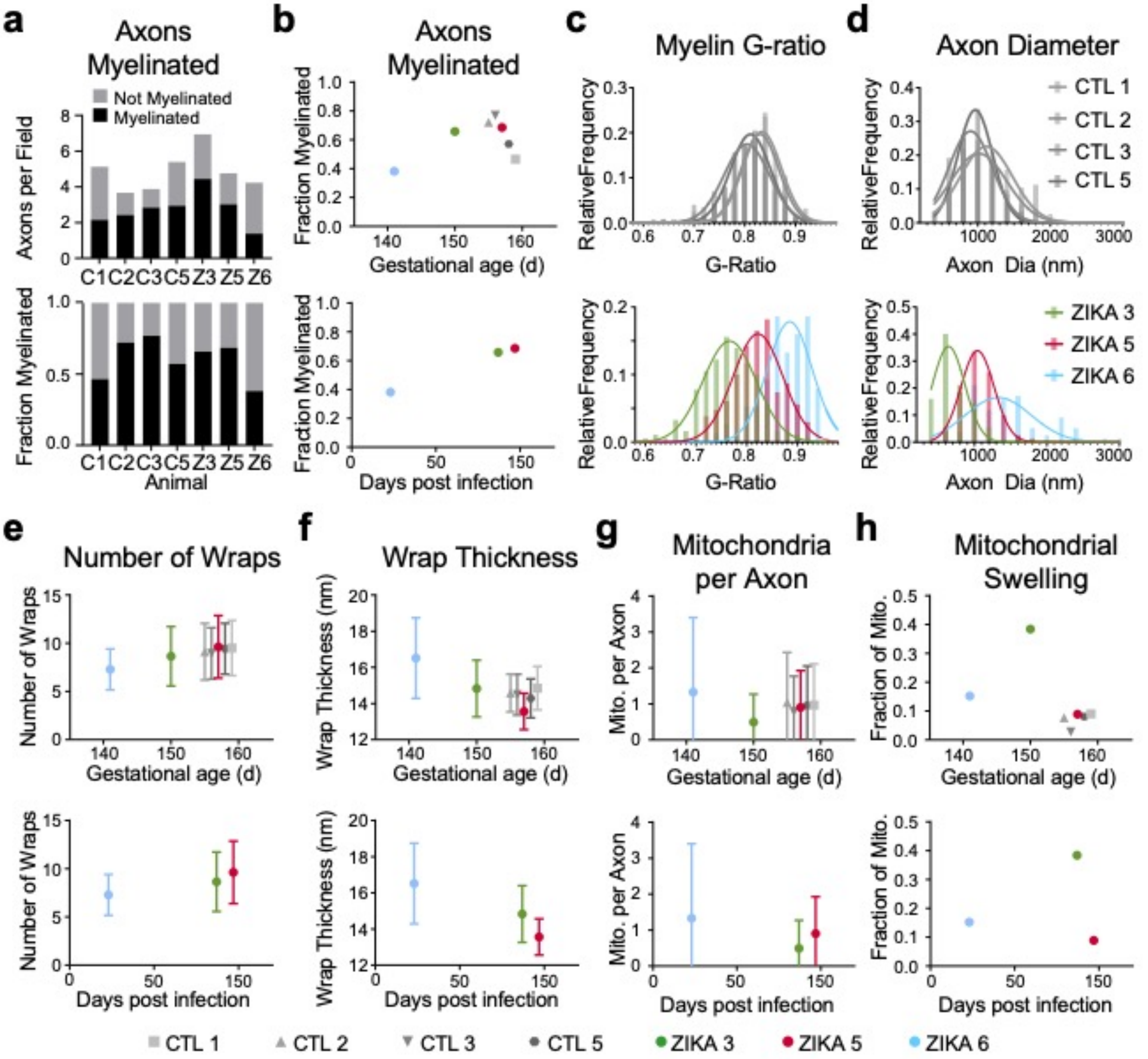


**Figure S7. Electron microscopy analysis of axon and myelin features in deep white matter of ZikV and control NHP fetal brains. a-b**, Quantification of the fraction of large-diameter (>250 nm) axons with **a**) mature myelin sheaths. Histograms representing **c**) myelin g-ratio and **d**) axon diameter across all analyzed axons for each animal in control (top) and ZikV-exposed (bottom) groups. Quantification of myelin features (e-f) and mitochondrial features (g-h) per axon in control and ZikV-exposed animals, plotted relative to gestational age (top row) or days post-ZikV inoculation (bottom row). Each colored point (graphs b, e-h) represents the average of all measurements for a single animal; error bars, when shown, indicate the mean and SEM of measurements across axons for each animal.

#### Figure S8

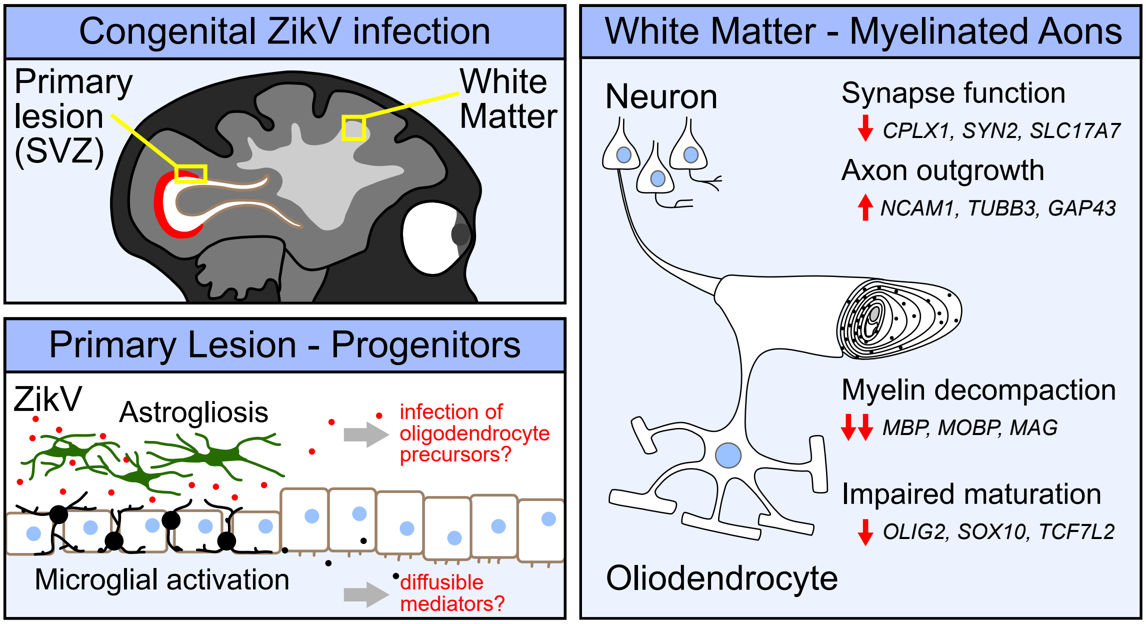


**Figure S8. Graphical abstract representing putative mechanisms for ZikV-induced myelin decompaction, including altered oligodendrocyte maturation, and neuronal function.** In a nonhuman primate model of non-microcephalic congenital Zika syndrome, we identified a primary lesion affecting the posterior periventricular area corresponding to the neural progenitor cell niche in the subventricular zone (SVZ), in which focal astrogliosis and reactive microglia were prominent features of pathology. In parietal cortex distal to this site, we found extensive perturbations of myelin with decompaction of the myelin sheath and loss of myelin basic protein expression. Spatially resolved transcriptional analysis identified ZikV-related changes in gene networks in the white matter with downregulation of oligodendrocyte functional genes, while in the grey matter genes for axon outgrowth were upregulated. We hypothesize congenital ZikV infection perturbs core oligodendrocyte transcriptional programs, either via direct infection of oligodendrocyte precursors or via diffusible mediators, leading to downregulation of genes necessary for maturation of oligodendrocyte lineage cells and strong downregulation of genes for myelin structural proteins. Loss of structural proteins such as myelin basic protein (MBP) leads to decompaction of myelin. In neurons, ZikV leads to loss of synaptic input, either because of decreased neurogenesis or because of dysfunctional electrical activity due to demyelination. In response, neurons increase expression of genes for axon outgrowth in order to make contact with new synaptic partners.
