## Supplemental Tables 1,2,10 for "Disruption of myelin structure and oligodendrocyte maturation in a pigtail macaque model of congenital Zika infection"

### SUPPLEMENTAL INFORMATION

#### Table of Contents

**Tables** ………………………………………………………………………………………………… 2-4

**Table S1.** Species, Sex, Age and Gestational Days of the Maternal Animals

**Table S2.** Species, Sex, Age and Weights of the Fetuses

**Table S3.** DSP differentially expressed gene matrix from **Fig. S2a**

**Table S4.** Representation gene set analysis from the heatmap shown in **Fig. S2a**

**Table S5.** GSEA results from DE genes by ROI, shown in **Fig. S2b**

**Table S6.** Ingenuity Pathway Upstream Regulator Analysis shown in **Fig. S2c**

**Table S7.** Bulk RNA-seq differential gene expression matrix from **Fig. S3c**

**Table S8.** Over Representation gene set analysis from the heatmap shown in **Fig. S3c**

**Table S9.** CIBERSORT analysis of bulk RNA-seq data shown in **Fig. S3d**

**Table S10.** Primary antibodies used for IHC and DSP analyses

**Supplemental Figure Legends**……..…………………………………………..…………... 5-10

#### TABLES

**Table S1. Species, Sex, Age and Gestational Days of the Maternal Animals**

| Study ID | Animal ID | Species | Sex | Age (y) | Gestational Age | | Days post-inoculation |
|---|---|---|---|---|---|---|---|
| | | | | | Inoculation | Delivery | |
| ZIKA1 | A10095 | *M. nemestrina* | F | 9.5 | 119 | 162 | 43 |
| ZIKA2 | F10094a | *M. nemestrina* | F | 5.9 | 82 | 159 | 77 |
| ZIKA3* | A10219a | *M. nemestrina* | F | 10.0 | 63 | 150 | 87 |
| ZIKA4* | T04054 | *M. nemestrina* | F | 12.8 | 60 | 142 | 82 |
| ZIKA5* | J07322 | *M. nemestrina* | F | 9.1 | 60 | 157 | 97 |
| ZIKA6 | A10008a | *M. nemestrina* | F | 12.2 | 121 | 141 | 20 |
| CTL1 | A07083 | *M. nemestrina* | F | 14.0 | 59 | 159 | 100 |
| CTL2 | M05062 | *M. nemestrina* | F | 12.0 | 63 | 155 | 92 |
| CTL3 | F09165a | *M. nemestrina* | F | 7.4 | 99 | 156 | 57 |
| CTL4 | A10183a | *M. nemestrina* | F | 7.8 | N/A | 156 | N/A |
| CTL5 | Z15006 | *M. nemestrina* | F | 5.2 | 138 | 158 | 20 |
| CTL6 | A10007 | *M. nemestrina* | F | 11.0 | 132 | 155 | 23 |

Abbreviations: F, Female; y, years (date of departure)

*Animals that received a mosquito salivary preparation inoculation and administration of DENV Ab, as previously described[1].

ZIKA refers to animal experiments with ZikV subcutaneous inoculation.

CTL, refers to animal experiments with media subcutaneous inoculation that underwent similar procedures as the ZikV animals.

N/A, not applicable

The average gestational age at delivery for pigtail macaques in the WaNPRC colony is 172 days gestation.

All animals were delivered by C-Section in the absence of labor.

**Table S2. Species, Sex, Age and Weights of the Fetuses**

| Study ID | Animal ID | Species | Sex | Age (d) | Body weight (kg) | Brain weight (kg) |
|---|---|---|---|---|---|---|
| ZIKA1 | Z16128 | *Macaca nemestrina* | M | 158 | 0.453 | N/A |
| ZIKA2 | Z16216 | *Macaca nemestrina* | F | 159 | 0.451 | N/A |
| ZIKA3 | Z16351 | *Macaca nemestrina* | F | 150 | 0.385 | 0.0514 |
| ZIKA4 | Z16354 | *Macaca nemestrina* | F | 142 | N/A | N/A |
| ZIKA5 | Z17003 | *Macaca nemestrina* | F | 157 | 0.453 | 0.0594 |
| ZIKA6 | Z19249 | *Macaca nemestrina* | F | 141 | 0.317 | 0.0472 |
| CTL1 | Z16296 | *Macaca nemestrina* | F | 159 | 0.432 | 0.0578 |
| CTL2 | Z17059 | *Macaca nemestrina* | F | 155 | 0.467 | 0.0563 |
| CTL3 | Z17004 | *Macaca nemestrina* | F | 157 | 0.352 | N/A |
| CTL4 | N/A | *Macaca nemestrina* | F | 158 | N/A | N/A |
| CTL5 | Z20054 | *Macaca nemestrina* | F | 158 | 0.499 | 0.0559 |
| CTL6 | Z21105 | *Macaca nemestrina* | M | 155 | 0.573 | 0.0601 |

Abbreviations: F, Female; M, Male; d, days; kg, kilogram

ZIKA, refers to animal experiments with ZikV subcutaneous inoculation.

CTL, refers to animal experiments with media subcutaneous inoculation that underwent similar procedures as the ZikV animals.

N/A, not available (not measured)

**Table S10. Primary antibodies used for IHC and DSP GeoMx analyses**

| Antibody | Source (cat #) | Species | Clone | Assay | Target | Dilution |
|---|---|---|---|---|---|---|
| GFAP | Dako (Z0224) | rabbit polyclonal | | IHC | astrocytes | 1:500 |
| AIF-1/Iba1 | Novus Biologicals, (MBP2-19019) | rabbit polyclonal | | IHC | microglia | 1:500 |
| Olig2 | Millipore (AB9610) | rabbit polyclonal | | IHC | oligodendrocytes | 1:500 |
| NeuN | Millipore (MAB377) | mouse monoclonal | Clone A60 | IHC | neurons | 1:500 |
| MBP | Abcam (ab7349) | rat monoclonal | Clone 12 | IHC | myelin sheath | 1:500 |
| RBFOX3 (NeuN) | Abcam (ab190195) | monoclonal | EPR12763 | DSP | neurons | 1:50 |
| Olig2 | Millipore (AB9610) | rabbit polyclonal | | DSP | oligodendrocytes | 1:100 |
| GFAP | Novus Biologicals (NBP-33184DL594) | mouse monoclonal | Clone GA-5 | DSP | astrocytes | 1:400 |
| STYO 83 | ThermoFisher (S11364) | | | DSP | nuclei | |
| Goat anti-Rb AF647 | ThermoFisher (A27040) | Goat | Superclonal | DSP | Rabbit IgG | |
